# NLRP1 inflammasome modulates senescence and senescence-associated secretory phenotype

**DOI:** 10.1101/2023.02.06.527254

**Authors:** Inés Muela-Zarzuela, Juan Miguel Suarez-Rivero, Andrea Gallardo-Orihuela, Chun Wan, Kumi Izawa, Marta de Gregorio-Procopio, Isabelle Coillin, Bernhard Ryffel, Jiro Kitaura, Alberto Sanz, Thomas von Zglinicki, Gabriel Mbalaviele, Mario D. Cordero

## Abstract

Senescence is a cellular aging-related process triggered by different stresses and characterized by the secretion of various inflammatory factors referred to as the senescence-associated secretory phenotype (SASP). Here, we present evidence that the inflammasome sensor, NLRP1, is a key mediator of senescence induced by irradiation both in vitro and in vivo. The NLRP1 inflammasome promotes senescence by regulating the expression of p16, p21, p53, and SASP in Gasdermin D (GSDMD)-dependent manner as these responses are reduced in conditions of NLRP1 insufficiency or GSDMD inhibition. Mechanistically, the NLRP1 inflammasome is activated downstream of the cytosolic DNA sensor cGMP-AMP (cGAMP) synthase (cGAS) in response to genomic damage. These findings provide a rationale for inhibiting the NLRP1 inflammasome-GSDMD axis to treat senescence-driven disorders.

## Introduction

Aging generates specific changes associated with a process called cellular senescence. This permanent state of cell cycle arrest promotes tissue remodelling during development but leads to the declined tissue regenerative potential and function after injury, and activates inflammation and tumorigenesis in aged organisms (1, 2). Senescence promotes the production of cytokines, chemokines, proteases, and growth factors some of which are known as senescence-associated secretory phenotype (SASP) (1). Recent studies demonstrates that chromatin is instrumental in regulating SASP and inflammation through the innate immune cyclic GMP-AMP synthase (cGAS)-stimulator of interferon genes (STING) which can be activated upon DNA sensing (3).

Inflammasomes are intracellular protein complexes involved in almost all human aging-associated complications such as cancer, cardiovascular, metabolic, and neurodegenerative diseases through the production of interleukin-1β (IL-1β) and IL-18 (4). These protein platforms comprise sensing proteins of the NOD-like receptor (NLR) family, the adaptor protein apoptosis-associated speck-like protein containing a CARD (ASC), and procaspase-1. Upon sensing of pathogen-associated molecular patterns (PAMPS) or damage-associated molecular patterns (DAMPS) (4), some NLRs such as NLRP3 and NLRP1 to some extent associate with ASC, a response that leads to the recruitment and activation of the cysteine protease, caspase-1 (4, 5). Active caspase-1 cleaves pro-IL-1β, pro-IL-18, and GSDMD, thereby facilatating the secretion of IL-1β and IL-18 through plasma membrane pores formed by the N-terminal fragments of GSDMD (4). These pores can also release IL-1α and cause pyroptosis (5). Despite scientific advances in the biology of the NLRP1 and NLRP3 inflammasomes, the role that these proteins play in senescence remain controversial (6-9).

Irradiation of cells or tissues is a widely used model of stress-induced senescence, which we used to determine the role of the NLRP1 and NLRP3 inflammasomes in this process. We found that irradiation induced the expression of NLRP1, NLRP3, and SASP. Notably, inhibition of the NLRP1 inflammasome but not NLRP3 inflammasome attenuated the expression of senescence markers, responses that were GSDMD- and cGAS-dependent.

## Results

### NLRP1 expression is stimulated during senescence

The ability of irradiation to cause cellular senescence provides an opportunity to study the role that NLRP1 and NLRP3 play in this model. Irradiation of human fibroblasts induced sustained expression of NLRP1, NLRP3, and senescence-associated proteins, p21 and IL-6, but transient expression of absent in melanoma 2 (AIM2) (Fig. 1A and fig. S1). Irradiation also stimulated the expression of IL-1β, IL-6, IL-8, and IL-18 as detected by ELISA (Fig. 1B and C). To reinforce the role that NLRP1 plays in SASP expression, we treated skin fibroblasts with Val-boroPro (VbP), a potent inducer of NLRP1 expression (10,11). VbP stimulated the expression of NLRP1, IL-1β, IL-18 and IL-6 (Fig. 1D and E). To confirm that SASP production is NLRP1-dependent, we determined the effects of siRNA-mediated NLRP1 knocked down on these responses. NLRP1 knock down inhibited the expression of p21, IL6, and IL-8 (Fig. 1F). NLRP3 Knock-down also downregulated IL-6 and IL-8 levels, but to a lesser extent compared to NLRP1 (Fig. S2A and B). These results suggest the NLRP1 inflammasome is the main regulator of SASP compared to the NLRP3 inflammasome.

**Figure 1.**
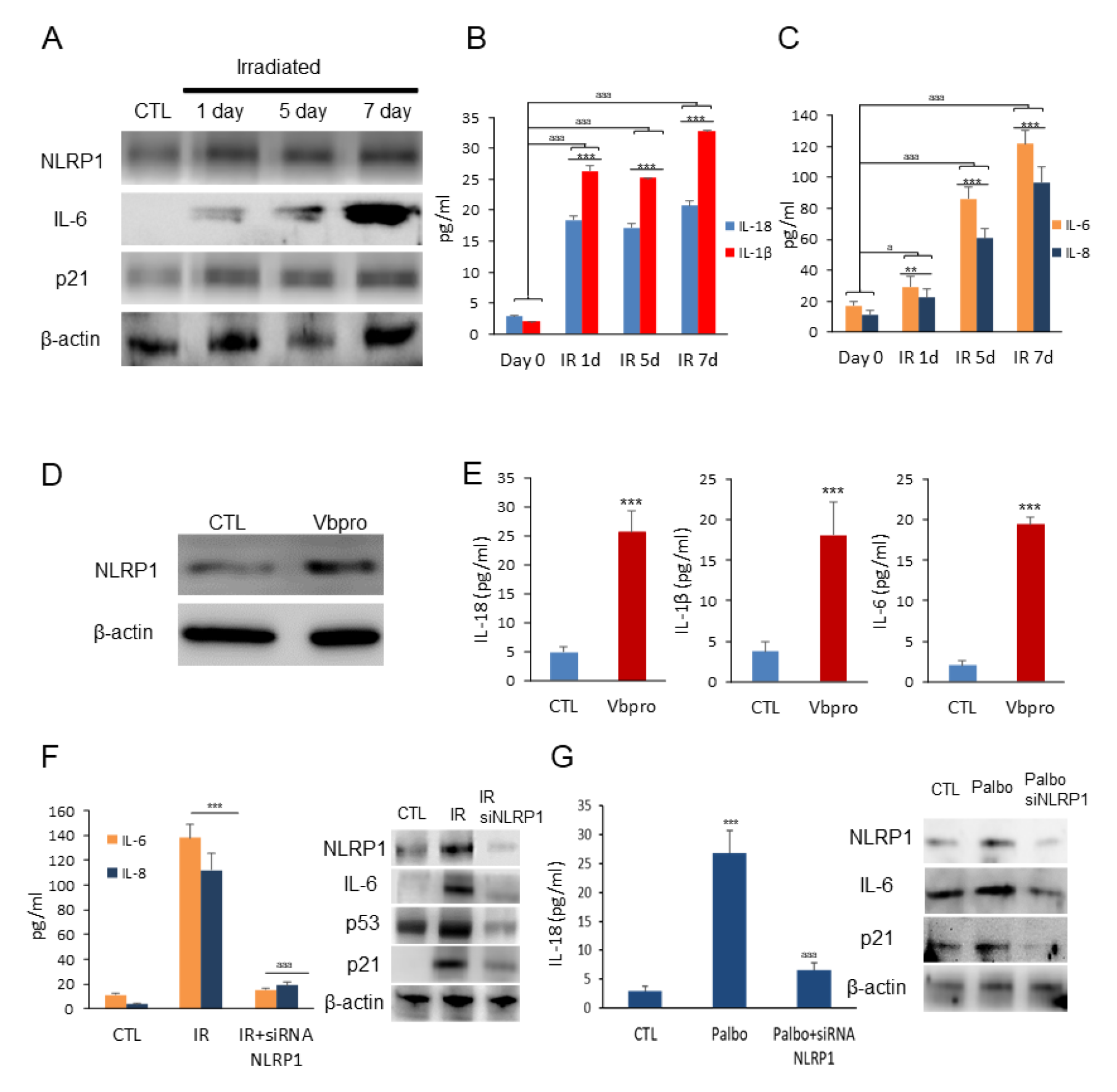
NLRP1 expression is associated with senescence. (A) Human fibroblasts were exposed to 20 Gy ionizing irradiation (IR). On day 1, 5 and 7, NLRP1 and senescence protein expression were analyzed by immunoblotting. (B and C) IL-1β, IL18, IL-6 and IL-8 were quantified by ELISA. All data are presented as means ± SEM, n = 3 independent experiments; **P < 0.005, ***P < 0.001 differences between time points after irradiation and day 0. (D and E) Human fibroblasts were treated wih Valboropro (VbP) to induce NLRP1 expression. After 24 h, NLRP1 and IL-6 protein expression were analyzed by immunoblotting and cytokines were analyzed by ELISA. (F and G) Human fibroblasts were irradiated (F) or stimulated with palbociclib (Palbo) (G) to induce two different senescence models. Then, cells were transfected with a non-targeting control siRNA (Control) or with siRNAs against NLRP1 (siNLRP1). Expression of NLRP1 and senescence-associated proteins p16, p21, p53 and IL-6 were assessed by immunoblotting and IL6, IL-8 or IL-18 were quantified by ELISA. All data are presented as means ± SEM, n = 3 independent experiments; **P < 0.005, ***P < 0.001 irradiated *vs* control. ^aaa^P < 0.001, IR+siRNA *vs* IR cells.

To determine whether the activation of NLRP1 is a common feature of different senescence inducers, we exposed human fibroblasts to palbociclib (PD-0332991), a CDK4 inhibitor, known to mimic the effect of p16^Ink4a^ (12-14). Palbociclib induced the expression of IL-18, NLRP1, IL-6, and p21, responses that were reduced in cells in which NLRP1 levels were down regulated by siNLRP1 (Fig. 1G). Thus, the expression of SASP is stimulated by two different activators of the NLRP1 inflammasome.

### NLRP1 inflammasome is necessary to induce paracrine bystander senescence

The NLRP1 inflammasome has been described as a sensor of various types of perturbations, modulting inflammation by mediating the secretion of pro-inflammatory cytokines, IL-18 and IL-1β (10,15). We hypothesized that the NLRP1 inflammasome promotes senescence by regulating SASP. To test this hypothesis, we exposed human skin fibroblasts to conditioned medium (CM) from human skin fibroblasts exposed to irradiation (supernatants of the cells cultured after 7 days of irradiation). We found that CM induced SASP responses in skin fibroblasts that were attenuated in cells transfected with siNLRP1 (Fig. 2A). CM from irradiated cells also impaired cell growth and promoted senescence as indicated by increased SA-Gal activity, IL-6 and IL-8 secretion, outcomes that were also decreased in siNLRP1-exposed cells (Fig. 2B-D).

**Figure 2.**
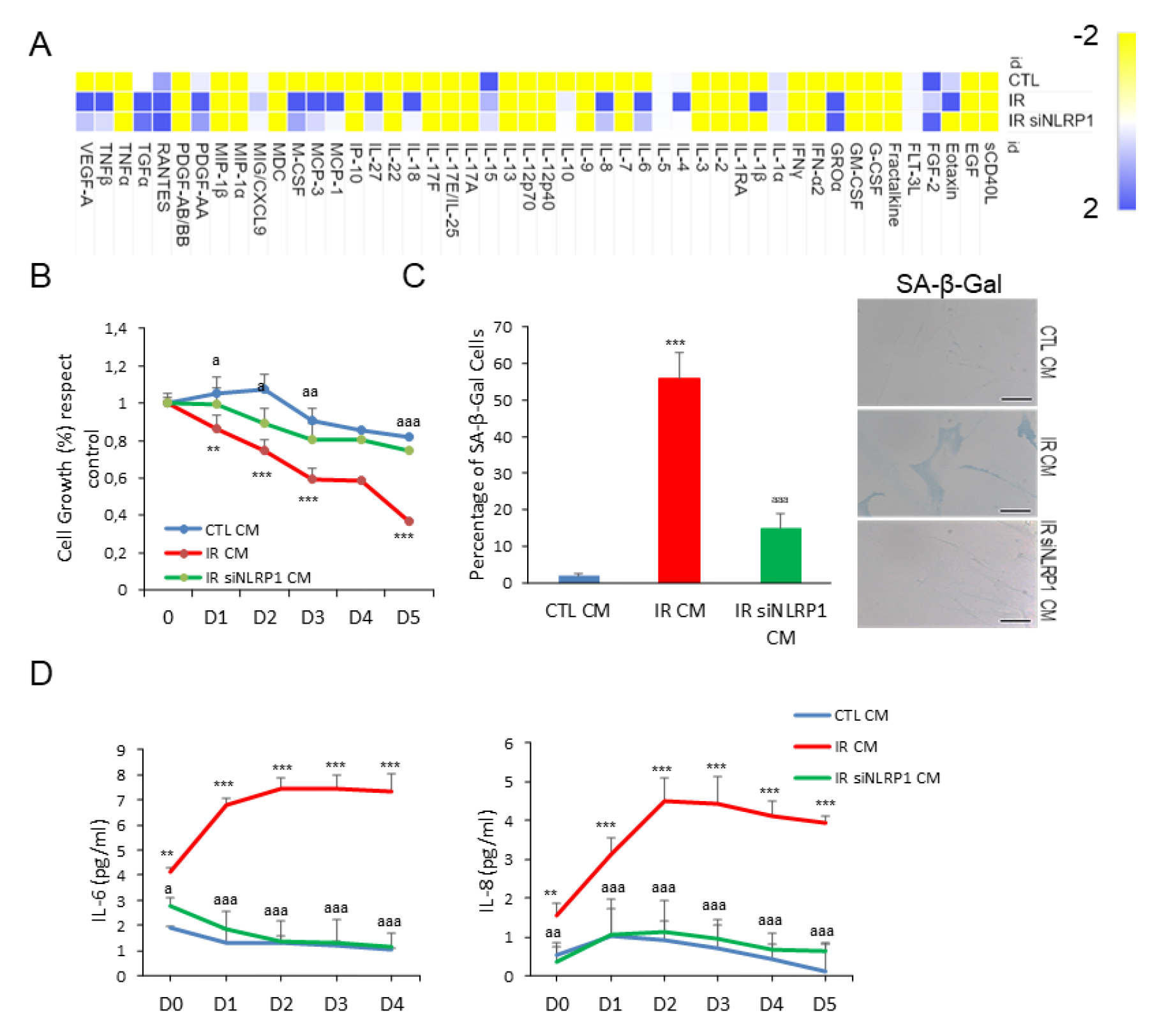
NLRP1 is necessary for paracrine senescence. (A) Heat map depicting expression of 48 human cytokines in the medium at 7 days following culture from IR and IR+siRNA NLRP1. (B) Effect of conditioned medium (CM) in human fibroblast growth. CM was collected from control, IR or IR+siNLRP1 cells. Percentage of cell growth was determined over 5 days. Image show cell populations stained with DAPI. (C) The induction of SA-β-Gal activity by CM was determined by microscopy. (D) IL-6 and IL-8 cytokine release to the medium after culture with CM from control, IR or IR+siNLRP1 cells. All data are presented as means ± SEM, n = 3 independent experiments; **P < 0.005, ***P < 0.001 irradiated *vs* control. ^a^P < 0.05, ^aa^P < 0.005, ^aaa^P < 0.001, IR+siRNA *vs* IR cells.

### NLRP1 deletion reduces senescence in vivo

To determine whether NLRP1 regulation of senescence occurs in vivo, we exposed wild-type (WT) mice and mice lacking all three alleles of NLRP1 (*Nlrp1a^−/−^Nlrp1b^−/−^Nlrp1c^−/−^*, referred to as *Nlrp1^−/−^*) to We hypothesized that the NLRP1 inflammasome promotes senescence by regulating SASP. To test this hypothesis, we exposed human skin fibroblasts to conditioned medium (CM) from human skin fibroblasts exposed to irradiation (supernatants of the cells cultured after 7 days of irradiation). WT mice also exhibited increased levels of IL-6, p16, p21, and several inflammatory mediators in the liver, outcomes that were decreased in *Nlrp1^−/−^* mice (Fig. 3D-F). To confirm the modulatory effect of NLRP1 on SASP, human fibroblasts were exposed to serum from irradiated mice. WT serum induced higher IL-6 levels compared to *Nlrp1^−/−^* serum (Fig. 3G). Histologic analysis revealed that irradiation damaged WT livers evidenced by sinusoidal dilatation and oedema (Fig. 3H) and increased nuclear p16^Ink4a^ expression (Fig. 3I), events that were prevented by NLRP1 deletion.

**Figure 3.**
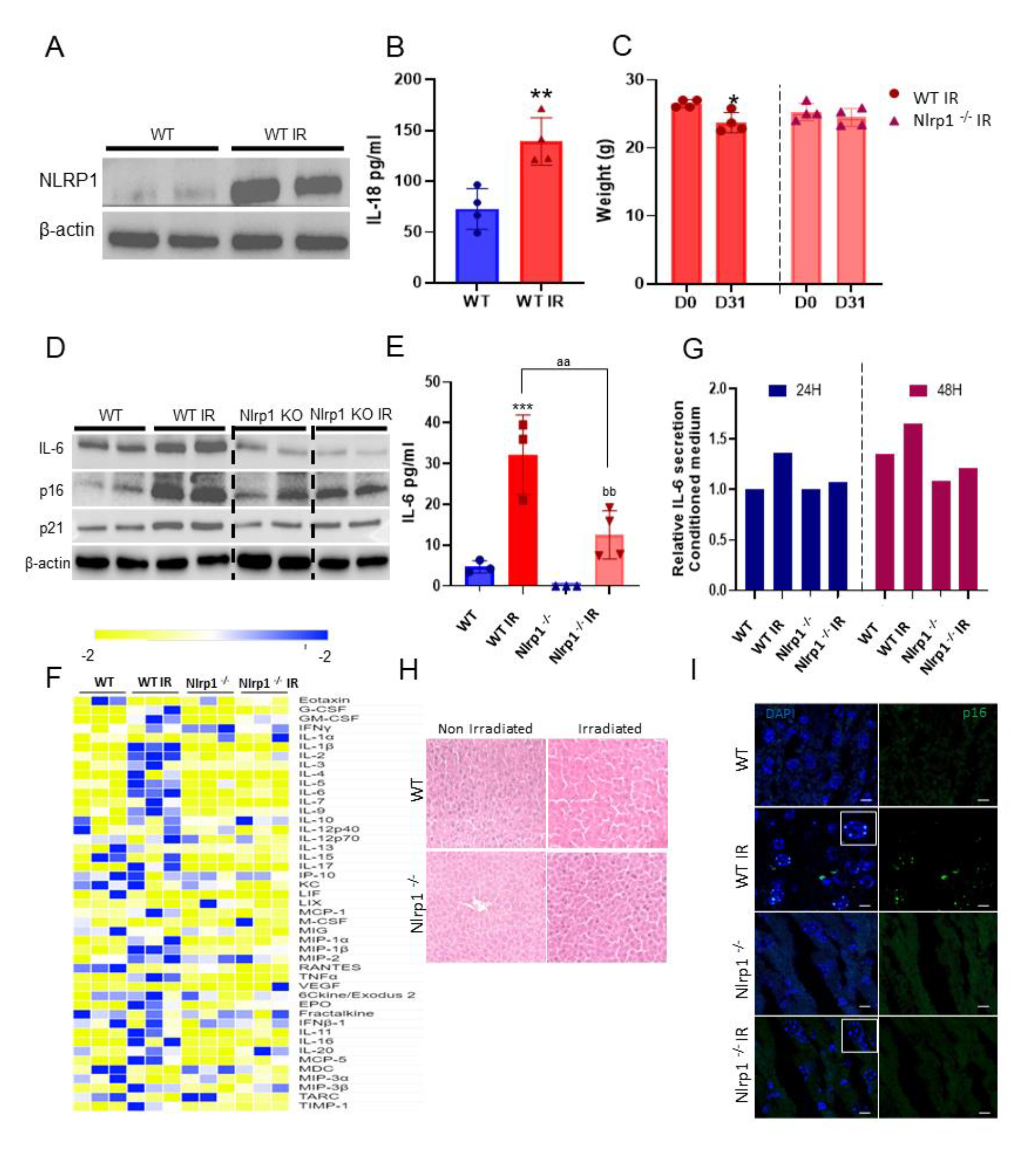
NLRP1 contributes to cellular senescence in vivo. (A) NLRP1 protein expression in liver from WT mice at one month after IR. (B) Serum levels of IL-18 after IR. (C) Effect of IR on the bodyweight of WT and NLRP1 knockout (KO) mice. (D) Protein expression in liver from IR and non-IR WT and NLRP1 KO mice of senescent markers (IL-6, p16 and p21). (E) Serum levels of IL-6 in serum from IR and non-IR WT and NLRP1 KO mice. (F) Heat map depicting expression of 44 mouse cytokines in serum at 5 weeks after IR of WT and NLRP1 KO mice. n = 3 mice per group. (G) IL-6 releases from healthy fibroblasts was assessed after 24 and 48 hr of incubation with media containing serum from IR and non-IR WT and NLRP1 KO mice. (H) Representative liver section stainings of hematoxylin and eosin (H&E). (I) Representative liver section immunostainings of p16. All data are presented as means ± SEM, n = 4-6 mice per group; *P < 0.05, **P < 0.005, ***P < 0.001 irradiated *vs* control. ^aa^P < 0.005 IR WT *vs* IR KO mice; ^bb^P < 0.005 IR KO *vs* IR KO mice.

We also determined NLRP3 actions on SASP in vivo. Unlike *Nlrp1^−/−^* mice, *Nlrp3 ^−/−^* mice showed significant weight loss after irradiation similar to WT mice (Fig. S3A). Irradiation also induced senescence and SASP markers p16, p21 and IL-6 to a similar extent in WT and *Nlrp3 ^−/−^* mice (Fig S3B-D). Together, these data indicate that NLRP1 but not NLRP3 control senescence and SASP in vivo.

### NLRP1 controls senescence in human tissues

To determine the relevance of NLRP1-mediated senescence in humans, we first examined p16^Ink4a^ expression in liver of a patient with NLRP1-dependent inflammasome hyperactivation caused by *de novo* c.3641C>T (p.P1214L) mutation in *NLRP1* (16). The patient showed clinical features associated with hyperkeratosis, liver cirrhosis and increased serum IL-18 but not of IL-1β. Immortalized hepatocytes from the patient released higher and lower levels of IL18 and IL1β, respectively (16). Accordingly, we found hyper-expression of p16^Ink4a^ in liver sections of this patient compared to a control liver samples (Fig. 4A), findings that were consistent with the previously reported association of senescence with liver steatosis and cirrhosis (17-20).

**Figure 4.**
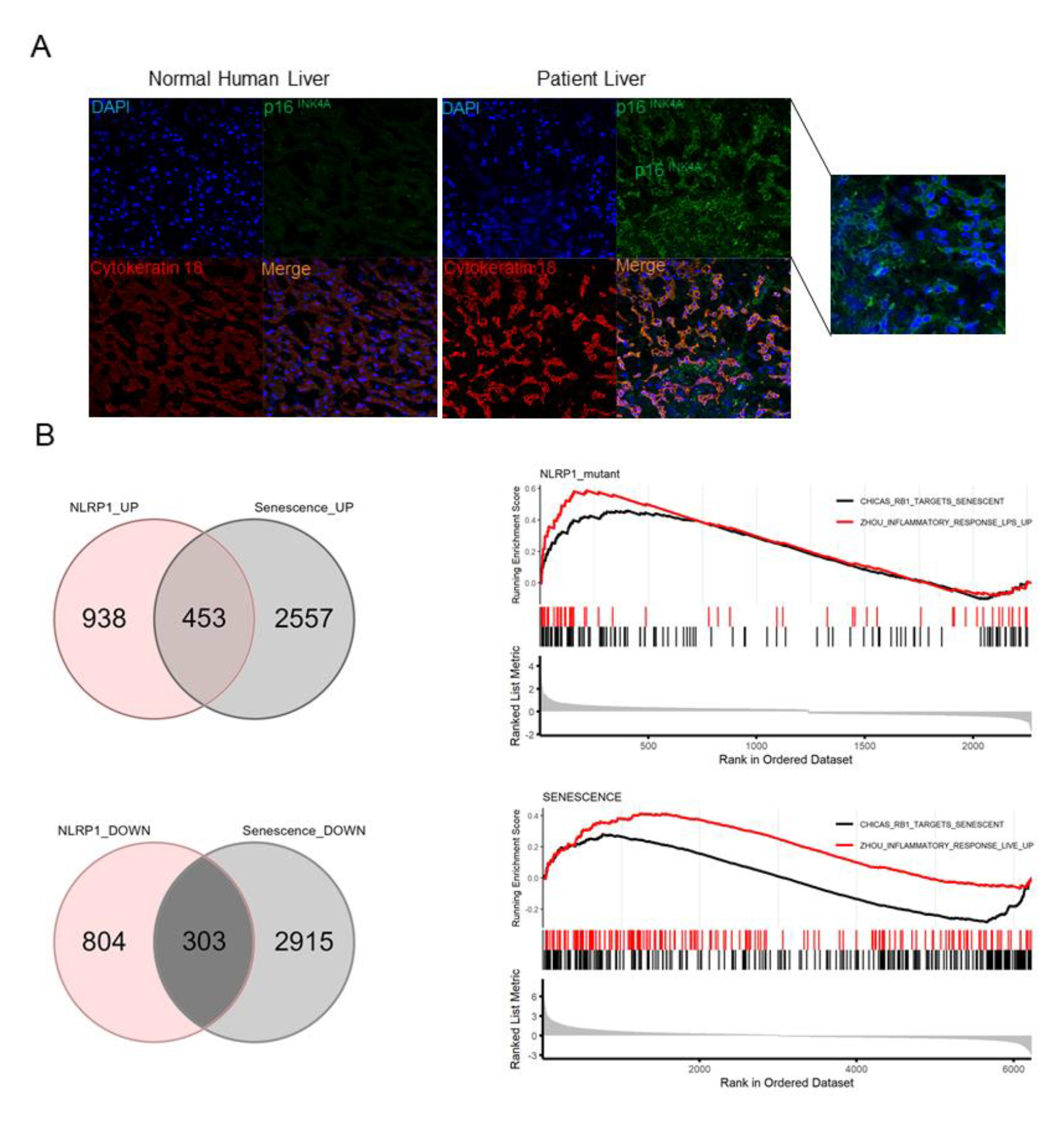
NLRP1 is associated with senescence in human. (A) p16 immunostaining in liver section from a patient with a NLRP1 gain of function mutation. (B) Venn Diagrams showing genes that are changed in both selected datasets (left), and genes that are upregulated (middle) or downregulated (right). GSEA analysis showing two gene datasets enrichered for inflammatory response and targets of senescences genes is shown below.

Finally, we compared the transcriptomes of keratinocytes from patients with germline mutations in *NLRP1* (21) *versus* a model of oncogene-induced senescence of these cells. We found that 756 transcripts were similar between both datasets, with 453 and 303 transcripts being up-and down-regulated, respectively. Furthermore, GSEA analysis revealed significant up-regulation of two sets of genes involved in the inflammatory responses and senescence (Fig. 4B). These results suggest that activation of the NLRP1 inflammasome triggers a transcriptomic reprogramming that is similar to the one occuring during senescence in humans.

### NLRP1 is activated by damaged DNA

DNA damage is commonly associated with senescence upon exposure to stressors such as ROS, radiotherapy, and chemotherapy (13); this insult is detected by various sensors, including cGAS (16). NLRP1 is known for sensing double-stranded (ds) RNA but not long dsDNA molecules (15, 16, 21). Given the role of NLRP1 as mediator of senescence and SASP as outlined above, we hypothesized that the NLRP1 inflammasome might be activated by damaged DNA. Therefore, we analyzed NLRP1 inflammasome-dependent responses upon exposure to human genomic DNA from irradiated cells (IR gDNA). We also used poly(deoxyadenylic-deoxythymidylic) acid (poly dA:dT) to activate the AIM2 inflammasome (22). IR gDNA increased NLRP1 and cGAS protein expression compared to non-IR gDNA (Fig. 5A), responses that correlated with IL-6 and IL-18 release (Fig. 5B). Moreover, poly(dA:dT) and irradiated Poly dA:dT oligonucleotide induced a significant overexpression of cGAS with IL-6 release but failed to induce NLRP1 expression or IL-18 release (Fig. 5C and D). These results suggest that NLRP1 expression is increased in response to DNA damage.

**Figure 5.**
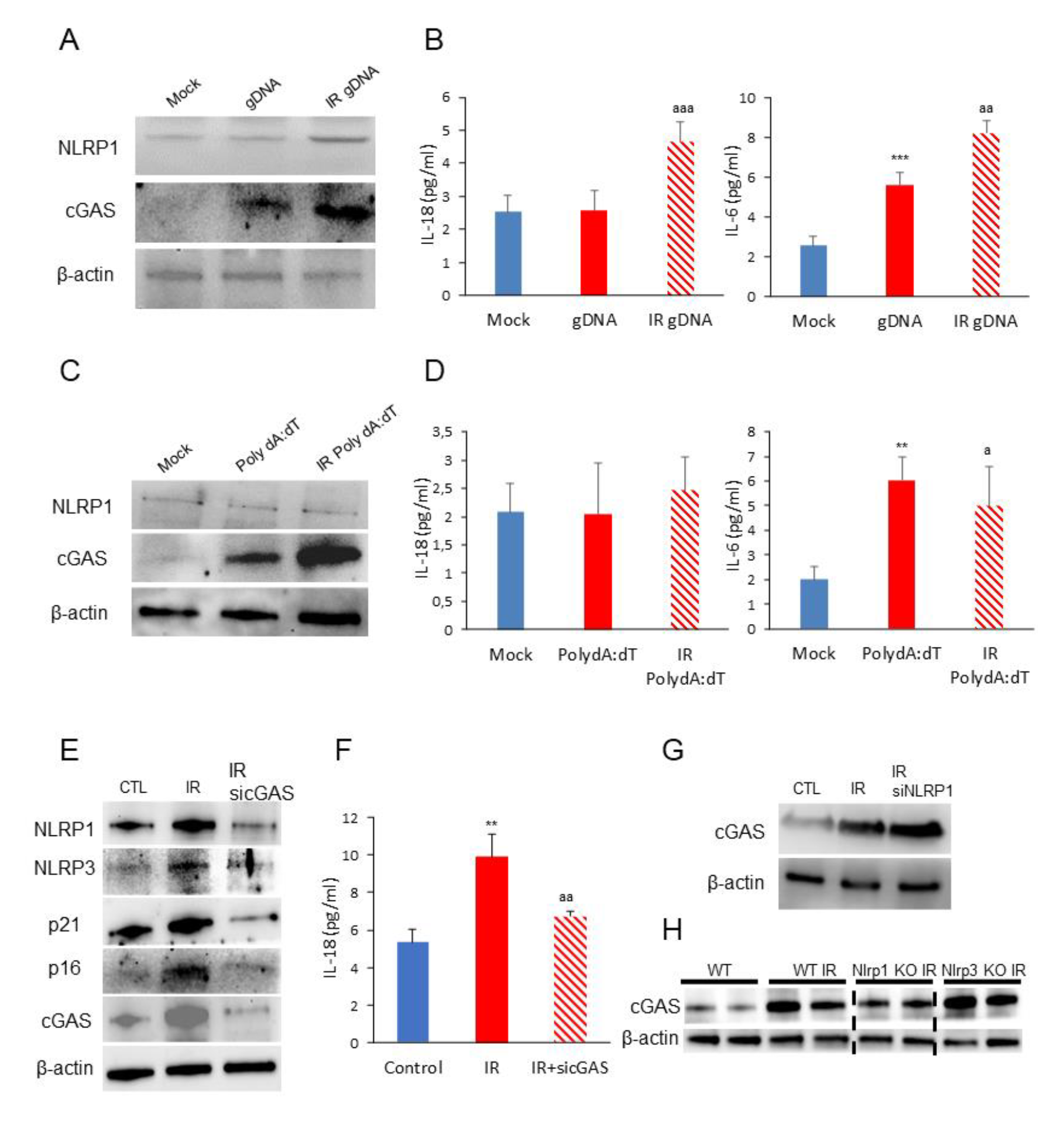
NLRP1 senses DNA damage mediated by cGAS activation. (A) Protein expression levels of NLRP1 and cGAS after 24 hr exposition to gDNA from non-irradiated and irradiated cells. (B) IL-6 and IL-18 release to the medium after the same experimental condition. Levels were determined by ELISA assay. (C and D) Protein expression levels of NLRP1 and cGAS and IL-6 and IL-18 release after 24 hr exposition to a non-irradiated or irradiated synthetic double-stranded DNA sequence, poly(dA-dT), Cytokine levels were determined by ELISA assay. Cytokine levels were determined by ELISA assay. All data are presented as means ± SEM, n = 3 independent experiments; **P < 0.005, ***P < 0.001 gDNA *vs* control cells. ^a^P < 0.05, ^aa^P < 0.005, ^aaa^P < 0.001, IR gDNA *vs* control cells. (E) Human fibroblasts were irradiated to induces senescence. Then, cells were transfected with a non-targeting control siRNA (Control) or with siRNAs against cGAS (sicGAS). Expression of NLRP1, NLRP3, IL-1β, cGAS and senescent protein p16, p21 was assessed by immunoblotting. (F) IL-18 release of non-irradiated, irradiated and irradiated and transfected with siRNAs against cGAS. Cytokine levels were determined by ELISA assay. All data are presented as means ± SEM, n = 3 independent experiments; **P < 0.005, IR *vs* contro cells. ^aa^P < 0.005, IR *vs* sicGAS.

To investigate whether NLRP1 expression depends on cGAS modulation, we knocked down cGAS with siRNAs in fibroblasts, which were then subjected to irradiation. Down regulation of cGAS resulted in decreased expression of not only p21 and p16 but also NLRP1, NLRP3, IL-1β, and IL-18 (Fig. 5E and F), suggesting that NLRP1 might modulate senescence downstream of cGAS. This view was supported by the fact that similar to NLRP1, cGAS expression was stimulated by irradiation (Fig. 5G and H). We noticed that lack of NLRP1 and NLRP3 led to increased cGAS abundancy (Fig. 5G), suggesting a compensatory effect on cGAS in the absence of NLRP1, findings that are consistent with the previously reported modulation of senescence and NLRP3 by cGAS pathway (13,16,23,24). These observations strengthen our view that NLRP1 expression is positively regulated by cGAS.

### NLRP1 controls SASP release through GSDMD pores

GSDMD pores release not only IL-1β and IL-18 (14), but also IL-33, another SASP member (25). Irradiation induced GSDMD cleavage in human fibroblasts, a response that we reduced by necrosulfonamide (NSA), an inhibitor of GSDMD (26). Irradiation-increased GSDMD expression and cleavage, responses that correlated with increased IL- 6 expression (Fig. 6A). NSA treatment reduced irradiation-induced GSDMD cleavage and cellular IL-6 abundance (Fig. 6A). Accordingly, NSA treatment completely blocked IL-6 and IL-18 secretion (Fig. 6B-C). Finally, we explored the role of GSDMD in SASP release *in vivo* by exposing Gsdmd KO (*Gsdmd^−/−^*) mice to total body irradiation and assessing a panel of SASP components in serum by cytokine array. Irradiation increased serum levels of various inflammatory factors, including SASP in WT mice, responses that were largely abrogated in *Gsdmd^−/−^* sera (Figs. 6D). These results suggest that the absence of GSDMD pores blocks the release of SASP factors in response to irradiation.

**Figure 6.**
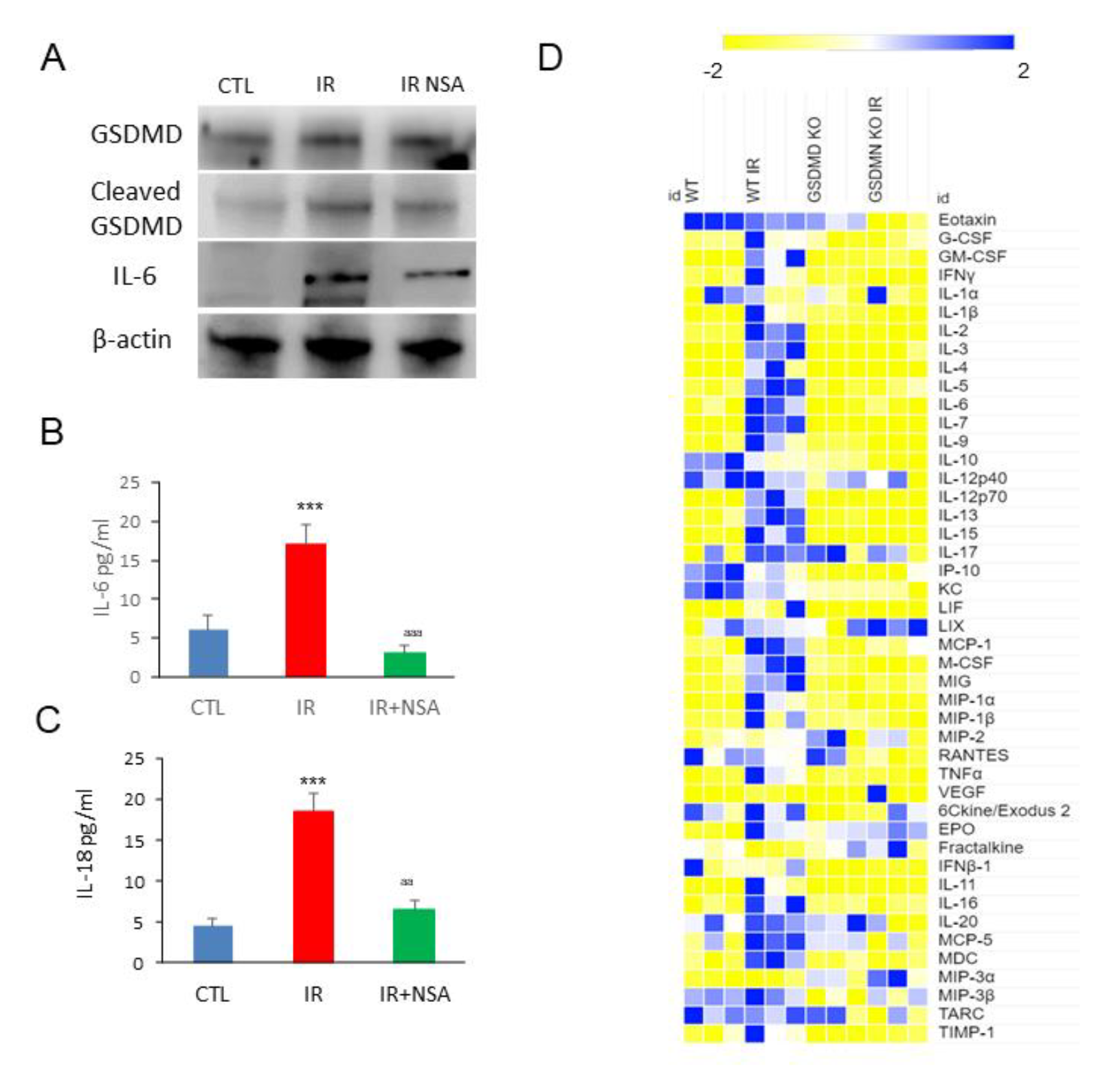
Gasdermin D mediates SASP release. (A) Protein expression levels of GSDMD, its cleaved form and of IL-6 in fibroblasts which were irradiated (IR) and treated with necrosulfonamyde (NSA) for 5 days after IR. (B and C) IL-6 and IL-18 release determined by ELISA assay. Data are presented as means ± SEM, n = 3 independent experiments; ***P < 0.001 irradiated vs control. ^aa^P < 0.005 IR cells vs IR NSA cells. (D) Heat map depicting expression of 44 mouse cytokines in serum at 5 weeks after IR of WT and GSDMD KO mice n = 3 mice per group.

## Discussion

We demonstrated that NLRP1 inflammasome acted downstream of cGAS to regulate senescence through GSDMD-dependent SASP secretion in various *in vitro* and *in vivo* models. cGAS recognizes diverse DNA species, including self-DNA (27,28), and its interactions with NLRP1 during senescence has not been reported. Our data revealed that the effects of the main inducers of cGAS-mediated SASP and senescence such as irradiation, palbociclib, or damaged DNA were regulated by the NLRP1 inflammasome and occurs through GSDMD pores. Recent works showing that human NLRP1 senses ultraviolet B (UVB) damage in skin are consistent with our observations since UVB also induces senescence (29-31). Our findings were also consistent with the association of NLRP1 overexpression or gain-of-function mutations with the prevalence of skin diseases such as psoriasis, vitiligo, atopic dermatitis or hyperkeratosis (32) where senescence had been widely described (33,34).

Our study showed increased levels of the senescence marker, p16^Ink4a^, in the liver of a patient with a *NLRP1* gain-of-function mutation. Furthermore, the transcriptome of NLRP1 gain-of-function mutations in human cells showed a significant inflammatory phenotype. This phenotype was associated with upregulation of stress-responsive secreted factors and known pro-inflammatory cytokines associated with SASP (35). The DPP8/DPP9 inhibitor Talablostat, also named PT-100 (Vbp), has been shown to induce human and rodent NLRP1 inflammasomes activation, and to stimulate the expression of SASP (36-39). Similarly, Maver et al., 2017 described homozygous missense variant in NLRP1 (Gly587Ser) in a family with multiple sclerosis with high IL-1β levels associated with several senescence and aging-associated genes such as NFKB, JNK and p38 pathways (35). Furthermore, Zhong et al., 2016 reported that germline NLRP1 mutations cause skin inflammation and cancer susceptibility (21). Collectively, our results unravel novel pathophysiological mechanisms of the NLRP1 inflammasome.

The NLRP1 inflammasome, like the NLRP3 inflammasome, has been associated with different age-related diseases (7,8). For example, increased protein levels of NLRP1 and caspase-1 is shown in the cortex of mice during aging (40). In the same line of thoughts, NLRP1 has also been proposed to contribute to the effect of age on chronic stress-induced depressive-like behaviour in mice (41). Because senescence is a very significant part of aging, these findings implicate NLRP1 in age-related diseases. Indeed, a recent study shows that the NLRP1 inflammasome is involved in age-related neuronal damage in which senescence markers are also described (42). NLRP1-siRNA treatment reduces neuronal senescence and damage associated with decreased β-galactosidase and apoptosis (43).

In conclusion, our findings provide valuable new insights into the mechanism of SASP production during senescence. Although several NLRP3 inhibitors have been developed, specific NLRP1 inhibitors are not currently available. Our study proposes NLRP1 as a new potential target to control cGAS-dependent inflammation and senescence reponses that drive human diseases.

## Material and Methods

### Reagents

Monoclonal antibodies specific for NLRP1 (NBP1-54899), NLRP3 (NBP2-12446) and p21/CIP1/CDKN1A (NBP2-29463) were purchased from Novus Biologicals (Colorado, USA). Similarly, anti-Caspase 1, IL-1β (D3U3E), cGAS (#79978), p16 ^INK4A^ (mouse: ab189034 and human: 18769S) were obtained from Cell Signaling Technology (Beverly, MA, USA). Finally, IL-6 (sc-32296), p53 (sc-126) antibodies and DAPI were obtained from Santa Cruz Biotechnology (Santa Cruz, CA, USA). Goat Anti-Rabbit IgG H&L (HRP), goat Anti-Mouse IgG, H&L Chain Specific Peroxidase Conjugate, Poly(deoxyadenylic-thymidylic) acid sodium salt (Poly dA-dT), BSA and Triton X-100 were obtained from Merck (Darmstadt, Germany). Val-boroPro - Calbiochem 5314650001 was obtained from Merck (Darmstadt, Germany). Necrosulfonamide was obtained from Sigma-Aldrich (Saint Louis, USA). A cocktail of protease inhibitors (Complete™ Protease Inhibitor Cocktail) was purchased from Boehringer Mannheim (Indianapolis, IN). The Immun Star HRP substrate kit was obtained from Bio-Rad Laboratories Inc. (Hercules, CA). Secondary Alexa Fluor 488 Goat Anti-Rabbit Antibody was obtained from Thermo Fisher (MA, USA). Finally, siRNAs of control and cGAS (AM16708 129125), NLRP3(4392420 S41555) and NLRP1 (4392420 S22520) were obtained from Invitrogen (Eugene, OR, USA).

### Cell culture

Primary human fibroblasts (Thermo fisher; C0135C) were cultured in high glucose DMEM (Dulbecco’s modified media) (Gibco, Invitrogen, Eugene, OR, USA) supplemented with 10% fetal bovine serum (FBS) (Gibco, Invitrogen, Eugene, OR, USA) and antibiotics (Sigma Chemical Co., St. Louis, MO, USA). Cells were incubated at 37°C in a 5% CO2 atmosphere. Senescent cells were generated by X-ray irradiation. 6300 cells/cm2 were seeded 24 h prior to 20 Gy irradiation and were used for experiments 7 days later

### Conditioned medium

Irradiated (20Gy) or non-irradiated cells (2×10^6^) were seeded in a 10 cm dish and incubated for 7 days in DMEM with 0.5% FBS. After incubation, the conditioned medium (CM) was collected, centrifuged at 5,000g and filtered through a 0:2 µm pore filter. CM was mixed with DMEM 40% FBS in a proportion of 3 to 1 to generate CM containing 10% FBS.

### Immunofluorescence assay

Fibroblasts were grown on 1 mm width glass coverslips for 72 hours in high glucose DMEM medium containing 10% FBS and 1% antibiotics. They were washed twice with PBS, fixed in 3.8% paraformaldehyde for 15’ at room temperature, permeabilized with 0.1% Triton X-100 in PBS for 10’ and incubated in blocking buffer (BSA 1%, Triton X- 100 0.05% in PBS) for 30’. In the meantime, the primary antibody was diluted 1:100 in antibody buffer (BSA 0.5%, Triton X-100 0.05% in PBS). Fibroblasts were incubated overnight at 4°C with the primary antibody and subsequently washed twice with PBS. The secondary antibody was similarly diluted 1:400 in antibody buffer, but their incubation time on cells was reduced to 2 hours at room temperature. Coverslips were then washed twice with PBS, incubated for 5’ with PBS containing DAPI 1 µg/ml and washed again with PBS. Next, they were mounted on microscope slides using Vectashield Mounting Medium (Vector Laboratories, Burlingame, CA, USA, H1000).

### SA-β-Galactosidase assay

5 ×10^4^ IMR90 cells per well were seeded in 6-well plates. Four days later, cells were fixed with 0.5% glutaraldehyde (Sigma) in PBS for 10 min. Fixed cells were washed three times with PBS 1mM MgCl_2_ pH 5.7, before adding to each well 2mL of pre-warmed X-Gal staining solution (2mM MgCl_2_, 5mM K_4_Fe(CN)_6_ - 3H_2_O, 5mM K_3_Fe(CN)_6_, 1 mg/mL X-Gal solution ready to use (R0941, ThermoFisher) in PBS). Plates were incubated for 2–24 h at 37 °C, washed and imaged. SA-β-Gal activity positive and negative cells were quantified using FIJI/ImageJ.

### ELISA (Enzyme-linked immunosorbent assay)

IL-6, IL-8, IL-1β and IL-18 levels were assayed in supernatant by duplicate using commercial ELISA kits (Thermo Fisher Scientific, MA, USA).

### Cytokine array

Blood serum was collected from wild-type and NLRP1, NLRP3 and GSDMD ^−/−^ mice. In an *in vitro* model of senescence induced by X-ray irradiation (10 Gy) (with or without irradiation), cells were cultured in serum-free media for 24 hr and media were collected for analysis. Media and blood serum were analyzed for expression of several mouse cytokine and chemokines (MD44) or human cytokine and chemokines (HD48), respectively, using a Multiplexing LASER Bead Assay (Eve Technologies).

### siRNA transfection

Cells were seeded on 6-well plates until 75% confluence in 2ml DMEM high glucose medium (Cat. 10566016) supplemented with 10% FBS and 1% antibiotics. Transfection was performed according to the lipofectamine RNAiMAX reagent (Cat. 13778-075) protocol. Briefly, the siRNA-lipid complex was prepared in DMEM medium with 3% lipofectamine and 30 pmol of the correspondent siRNA, and incubated for 5 minutes at RT to form the silencing complex. Then, 250 µl of the siRNA-lipid complex were added to each well. After 72h, cells were treated and analyzed for the different conditions. Every reagent, including DMEM medium, was purchased from ThermoFisher (Waltham, MA, USA).

### DNA treatment

For DNA extraction, cells were seeded on T75 flasks until 80% confluence, then cells were irradiated (20 Gy irradiation and were used for experiments 7 days later) using X- RAD225, 225 kV. After one week, irradiated and control cells were scrapped off and centrifuged at 1.000g for 5’. DNA extraction was performed using 500 μl lysis buffer containing 100mM Tris HCl, 100mM EDTA, 100mM NaCl, SDS 1%, pH 7.5. Samples were incubated at 65°C for 30’, and 500 μl Phenol/Chloroform/Isoamyl Alcohol 25:24:1 was added. Samples were mixed by simple inversion and centrifuged at 5.000g for 5’. 300 μl DNA-containing top aqueous phase was retrieved. For DNA precipitation, samples were mixed with 200 μl 5M potassium acetate and 400 μl isopropanol. After 12.000g for 15’ centrifugation, the pellet was washed with 70% ethanol twice and dried for 1 hour. DNA pellet was resuspended in TE buffer (10mM Tris HCl, 1mM EDTA, pH 7.5) and quantified using a NanoDrop™ One/One from Thermo Fisher (Waltham, MA, USA).

Non-irradiated and irradiated DNA-treated cells were seeded on 6-well plates until 90% confluence in 2ml DMEM high glucose medium supplemented with 10% FBS and 1% antibiotics. Then, 1 µg/ml of control/ irradiated DNA was added for 8h/24h. Finally, cells were scrapped off and pelleted for further analysis.

Poly(dA-dT) DNA was transfected using Lipofectamine 2000 at 1 µg/ml concentration for 24h. Cells were scrapped off and pelleted for further analysis.

### Immunoblotting

Western blotting was performed using standard methods. After protein transfer, the membrane was incubated with various primary antibodies diluted 1:1000; the corresponding secondary antibodies were coupled to horseradish peroxidase at a 1:10000 dilution. Specific protein complexes were identified using the Immun Star HRP substrate kit (Biorad Laboratories Inc., Hercules, CA, USA).

### Ethical Statements

Animal studies were performed in accordance with the European Union guidelines (2010/63/EU) and the corresponding Spanish regulations for the use of laboratory animals in chronic experiments (RD 53/2013 on the care of experimental animals). All experiments were approved by the local institutional animal care committee.

### Animals

For all experiments, only male mice were used. WT C57/BL6/J, Nlrp1a^−/−^Nlrp1b^−/−^Nlrp1c^−/−^ (Nlrp1^−/−^) and NLRP3^−/−^ transgenic mice (C57BL/6J background), weighing 25-30 g, were maintained on a regular 12 h light/dark cycle. All groups had *ad libitum* access to their prescribed diet and water throughout the study. Body weight and food intake were monitored weekly. Animal rooms were maintained at 20– 22°C with 30–70% relative humidity.

### Irradiation

At 5–6 months of age, mice were sub-lethally irradiated (NDT 320 or X-RAD225, 225 kV) with a total of 12 Gy of X-ray irradiation (3 times 4Gy, with 2 days recovery between doses). Two days prior to irradiation (IR), and for 14 days post-IR, mice received 1% Baytril solution (Broad-spectrum antibiotic) in drinking water. At one month after IR, mice were sacrificed at the end of the study by cervical dislocation and tissues harvested, and stored in 4% paraformaldehyde for 24 hours for paraffin embedding, or frozen in liquid nitrogen. Blood samples were isolated by cardiac puncture.

### Histological study

After sacrifice of mice, livers were excised and immediately stored in 4% paraformaldehyde at room temperature for 24 hours for paraffin embedding after a brief rinse with PBS. The specimens were cut into 5-μm sections and stained with hematoxylin and eosin.

### Immunofluorescent Staining of Paraffin-embedded Sections

Paraffin sections were attached to superfrost plus slides (Menzel-Glaser, Braunschweig, Germany) at 60°C for 1 hour. Deparaffinization was performed by pure xylol washes 3 times for 10’ each. Slides were rehydrated by ethanol solutions immersion (from 100% to 70%) for 5’ each and rinsed with deionized water. For the heat antigen retrieval, slides were immersed in sodium citrate 10mM (unmasking solution) and microwaved at 800 W for 15’, then samples were kept at room temperature until cool down. Slides were rinsed with PBS 1×3 times, and then blocking solution (2% BSA, 0.05% Triton X-100 in PBS 1x) was applied for 1 hour. The samples were surrounded with a hydrophobic barrier using a barrier pen, and the primary antibody was applied at 1:100 concentration and diluted in a blocking solution overnight. The next day, slides were rinsed 3 times with PBS 1x, and the secondary antibody was applied at 1:400 concentration diluted in blocking solution for 2 hours. Again, slides were rinsed 3 times with PBS 1x, and DAPI staining (1ug/ml) was applied for 10’. Finally, samples were mounted with coverslips using Vectashield Mounting Medium (Vector Laboratories, Burlingame, CA, USA, H1000).

### Bioinformatics analysis

We used ARCHS4 (44) to download the samples of the two datasets analyzed referring to mutations in NLRP1 and a model of oncogene-induced senescence in keratinocytes, GSE85791 & GSE180361, respectively. ARCHS4 provides RNAseq data that are uniformly processed using the Kallisto aligner (45). Filtering and normalization were performed using the TMM method in EdgeR (3.40.1). Differentially expressed genes were obtained using the VOOM function from the Limma package (3.54.0). An FDR < 0.05 was used as a cut-off. Finally, differentially expressed genes from both datasets were individually selected for Gene Set Enrichment Analysis (GSEA) using GSEA from the GSEABase package (1.6.0), selecting hs_gsea_c2 as the geneset.

### Statistics

All data are expressed as means ± SEM. After evaluation of normality using Shapiro-Wilk test, statistical differences among the different groups were measured using either an unpaired Student t-test or 1-way analysis of variance (ANOVA) when appropriate with Tukeys post-hoc test. A P value of ≤0.05 was considered statistically significant. Statistical analyses were performed using Prism software version 5.0a (GraphPad, San Diego, CA). Asterisks in the figures represent the following: *: P ≤0.05; **: P ≤ 0.01; and ***: P ≤ 0.001.

## Acknowledgments

We thank Dr. Seth L. Master for providing the NLRP1 KO mice. This study was supported by PI21/01656 grant, Instituto de Salud Carlos III, Spain.

## Funding

MDC is supported by PI21/01656 grant from Instituto de Salud Carlos III, Spain and PRF 2021-78, Progeria Research Foundation. GM is supported by AR076758-01, R01AI161022, and R01AG071085 grants from NIH. BR and IC are supported by CNRS, University of Orleans, ‘Fondation pour la Recherche Médicale’ (EQU202003010405) and European funding in Region Centre-Val de Loire (FEDER N° EX010381). AS is supported by Wellcome Senior Research Fellowship (212241/A/18/Z) & BBSRC grants (BB/R008167/1 & BB/W006774/1).

## Conflict of interests

The authors declare that the research was conducted in the absence of any commercial or financial relationships that could be construed as a potential competing interest.

## Supplementary Data

**Supplementary Figure 1.**
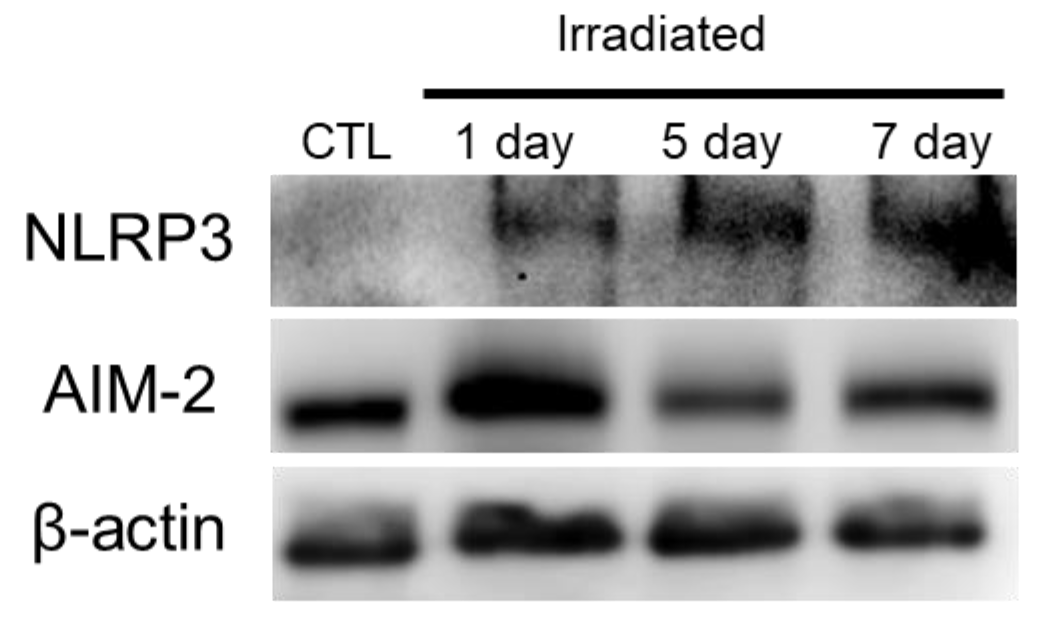
NLRP3 is expressed during senescence induced by irradiation. Human fibroblasts were exposed to 20 Gy ionizing irradiation. At day 1, 5 and 7 NLRP3 protein expression was analysed by immunoblotting after irradiation. n = 3 independent experiments.

**Supplementary Figure 2.**
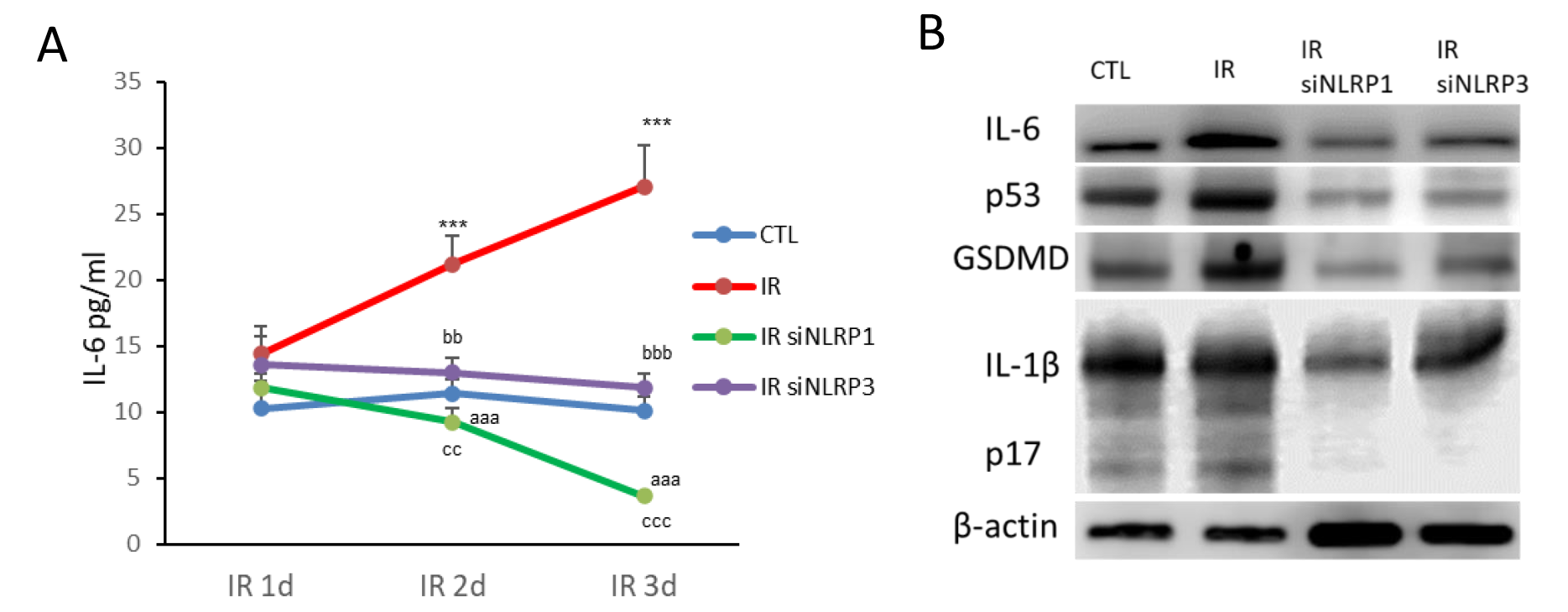
NLRP1 induces a more significant senescenct control than NLRP3. (A) IL-6 release from cell IR, IR+siNLRP1 or IR+siNLRP3 cells and (B) protein expression All data are presented as means ± SEM, n = 3 independent experiments; ***P < 0.001 irradiated *vs* control. ^aaa^P < 0.001, IR+siRNA NLRP1 *vs* IR cells; ^bb^P < 0.005, ^bbb^P < 0.001, IR+siRNA NLRP3 *vs* IR cells; ^cc^P < 0.005, ^ccc^P < 0.001, IR+siRNA NLRP1 *vs* IR+siRNA NLRP3 cells.

**Supplementary Figure 3.**
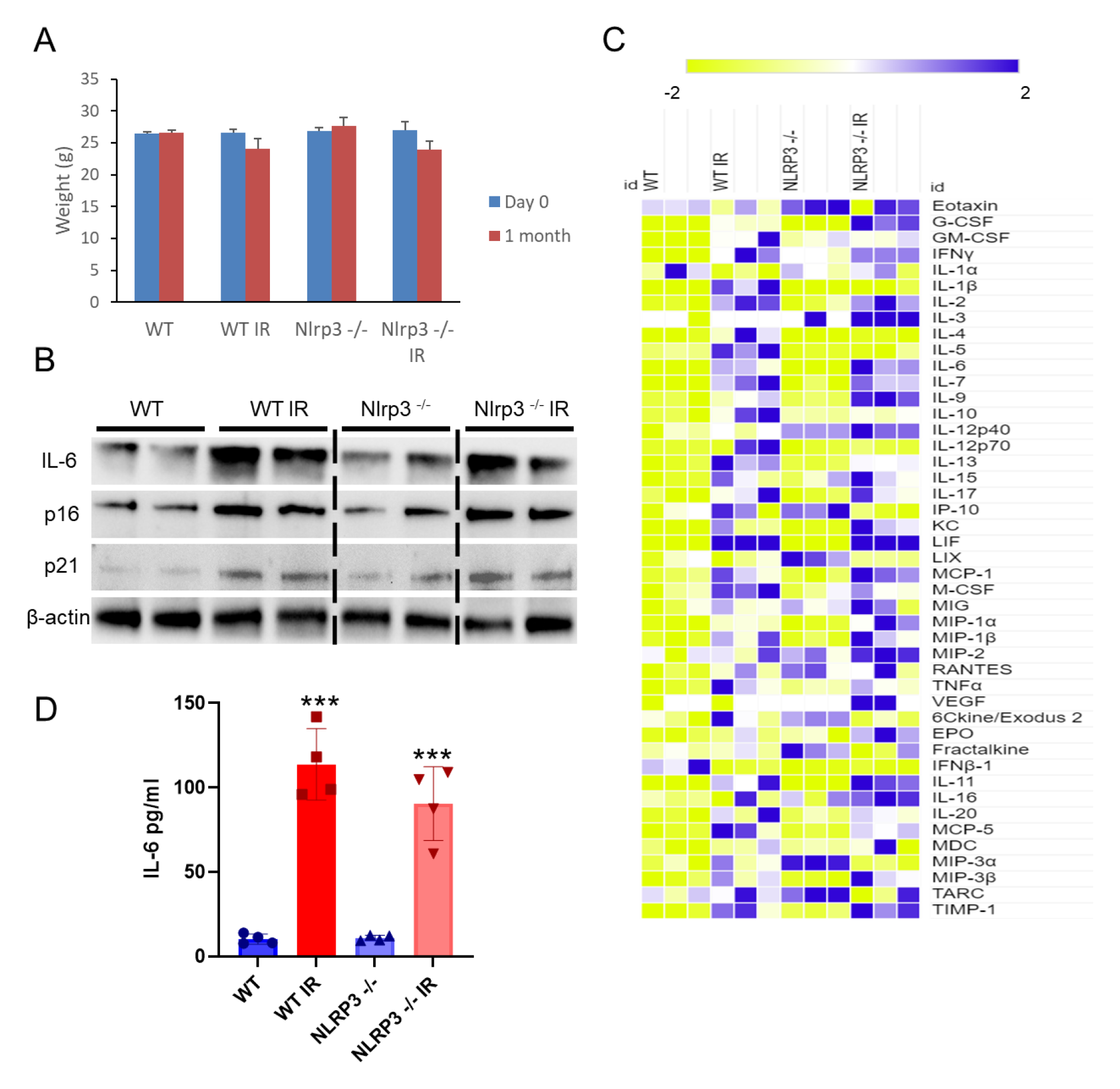
NLRP3 deletion does not protect of senescence and SASP. (A) Effect of IR on the bodyweight of WT and NLRP3 knockout (KO) mice. (B) Protein expression in livers from IR and non IR-WT and NLRP3 KO mice of senescence protein markers (IL-6, p16 and p21). (C) Heat map depicting expression of 44 mouse cytokines in serum at 5 weeks following from IR of WT and NLRP3 KO mice. (D) IL-6 release from healthy fibroblasts was assessed after 24 and 48 hr of incubation with media containing serum from IR and non IR WT and NLRP3 KO mice. All data are presented as means ± SEM, n = 3-6 mice per group; ***P < 0.001 irradiated *vs* control.

## Notes

### Competing Interest Statement

The authors have declared no competing interest.

### Summary of Updates

Figure 1 revised

